# Genomic Resources of Broomcorn Millet: Demonstration and Application of a High-throughput BAC Mapping Pipeline

**DOI:** 10.1101/2020.11.19.389643

**Authors:** Wei Xu, Mengjie Liang, Xue Yang, Hao Wang, Meizhong Luo

## Abstract

**Background:** With high-efficient water-use and drought tolerance, broomcorn millet has emerged as a candidate for food security. To promote its research process for molecular breeding and functional research, a comprehensive genome resource is of great importance.

**Results:** Herein, we constructed a BAC library for broomcorn millet, generated BAC end sequences based on the clone-array pooled shotgun sequencing strategy and Illumina sequencing technology, and integrated BAC clones into genome by a novel pipeline for BAC end profiling. The BAC library is consisted of 76,023 clones with an average insert length of 123.48 Kb, covering about 9.9-fold of the 850 Mb genome. Of 9,216 clones tested using our pipeline, 8,262 clones were mapped on the broomcorn millet cultivar longmi4 genome. These mapped clones covered 308 of the 829 gaps left by the genome. To our knowledge, this is the only BAC resource for broomcorn millet.

**Conclusions:** We constructed a high-quality BAC libraray for broomcorn millet and designed a novel pipeline for BAC end profiling. BAC clones can be browsed and obtained from our website (http://eightstarsbio.com/gresource/JBrowse-1.16.5/index.html). The high-quality BAC clones mapped on genome in this study will provide a powerful genomic resource for genome gap filling, complex segment sequencing, FISH, functional research, and genetic engineering of broomcorn millet.

## 1 Introduction

With the increased global water scarcity caused by climate change and population growth, it is of great importance to exploit the high-efficient water-use crop for the food security of the human in the future. Broomcorn millet (*Panicum miliaceum*), also known as proso millet, panic millet, and wild millet, is one of the traditional five-grain crops in the north of China [1]. It is a typical C4 plant with high photosynthetic efficiency. It also has a high-efficient utilization ratio of water resource and a capacity for drought resistance, exquisitely adapting to semi-drought or drought conditions [2]. Furthermore, its growing cycle, 60–90 days from sowing to maturity, is shorter than other cereals [3]. Broomcorn millet contains more protein than most grains, and a relatively balance array of trace elements and vitamins. More than 8700 accessions of broomcorn millet including landraces and cultivars have been conserved in the National Gene Bank of the Institute of Crop Science, Chinese Academy of Agricultural Sciences, thus providing an abundance of resources for genetic improvement of broomcorn millet.

At present, high-quality chromosome-scale genome assemblies of two allotetraploid (2n = 4x = 36) broomcorn millet varieties decoded by Chinese researchers are available [4, 5]. These genome assemblies provide the foundation for the molecular breeding of broomcorn millet. However, other gemonic resources are still required to complete and make the full use of the genome assemblies.

The tranditional bacterial artificial chromosome (BAC) libraries with genomic DNA inserts of 50 kbp – 300 kbp [6–8] are also important resources for genomic research. They provide natural DNA materials for a variety of experiments, such as intact gene cluster cloning, map-based cloning, whole genome sequencing, comparative genomics analysis and fluorescence *in situ* hybridization that aim to understand functional elements in the genome [9, 10]. The utility of the BAC libraries can be greatly enhanced by mapping the BAC clones on the genome assemblies.

Hence, we constructed a high-quality BAC library for broomcorn millet, developed a new pipeline for cost-effectively decoding BAC end seqeunces generated by clone-array pooled shotgun sequencing strategy (abbr. CAPSS) and Illumina sequencing technology, and mapped the BAC clones on the genome assemblies of the broomcorn millet [11].

## 2 Results

### 2.1 BAC library construction

A BAC library of broomcorn millet was constructed with the restriction enzyme *Hin*dIII using high-molecular-weight genomic DNA prepared from etiolated seedlings. In total, the library consists of 76,032 BAC clones, that were arrayed into 198 independent 384-well microtiter plates. Insert sizing of randomly picked clones showed that the majority of genomic BAC inserts fell into the length range of 97–145.5 kb with an average insert size of 132 kb (Fig. 1).

**Fig. 1.**
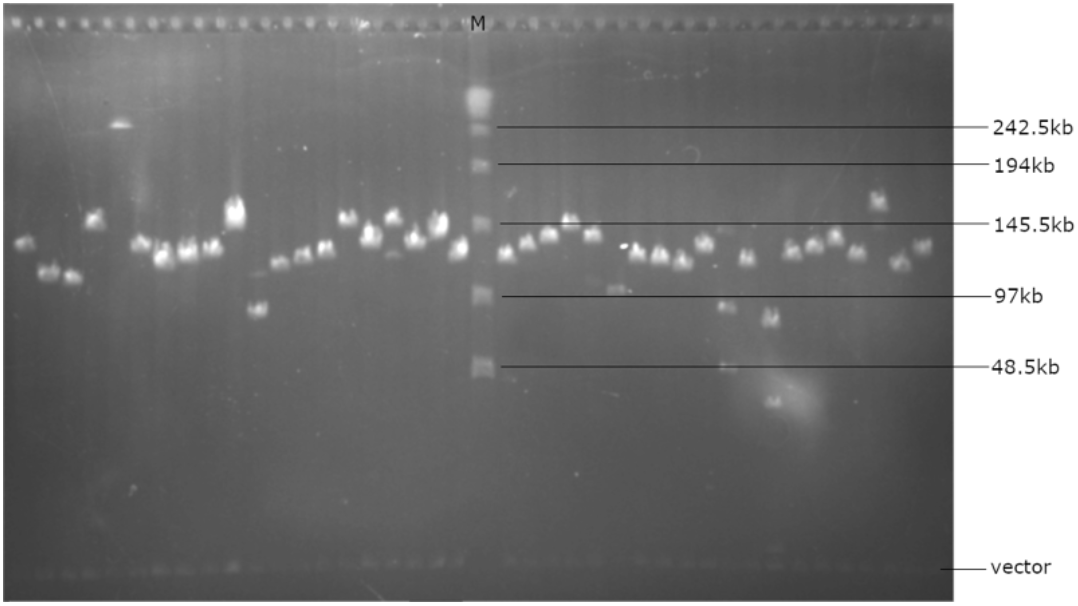
The insert sizes of randomly selected BAC clones determined by PFGE. The maker in the middle is *λ* DNA ladder.

### 2.2 Construction and sequencing of DNA pools

In order to obtain BAC end sequences with high efficiency and low cost, we designed and performed a pipeline based on the clone array pooled shotgun sequencing strategy (Fig. 2). Randomly selected 24 384-plates (each plate consists of 16 x 24 clones) from the broomcorn millet BAC library were arranged to a square superpool with 6 row plates and 4 column plates. Therefore, the superpool consisted of 96 row pools (6 plates x 16) and 96 column pools (4 plates x 24). These row pools and column pools were called as the secondary pools, and each secondary pool also consisted of 96 BAC clones. In total, one superpool consisted of 192 secondary pools and 9,216 individual BAC clones.

**Fig. 2.**
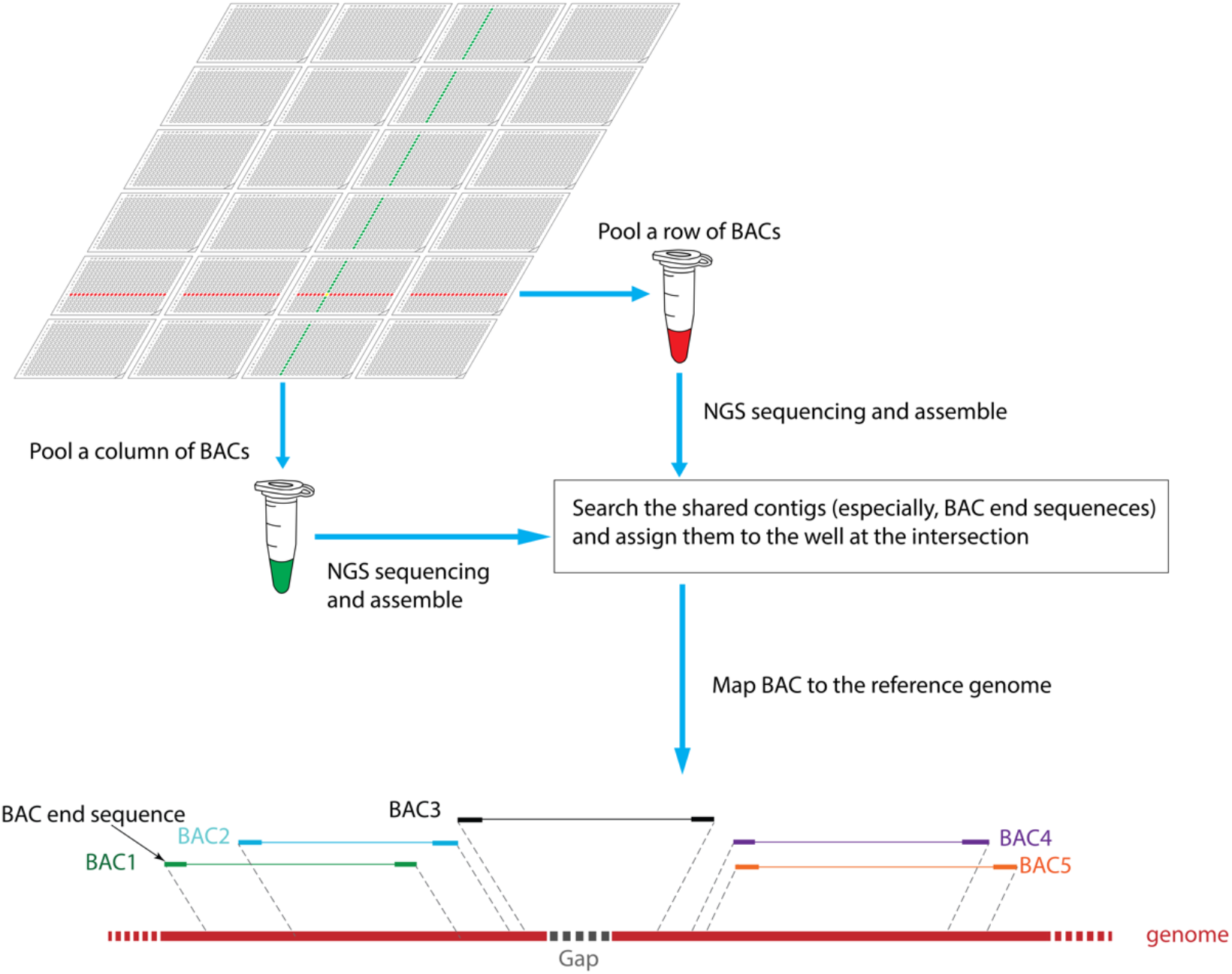
The strategy of BAC end sequence localization based on CAPSS. Twenty-four 384-plates are arranged to a square superpool, and then 96 row pools and 96 column pools are prepared and sequenced by NGS platform. The sequences of each pool are assembled into contigs. If a contig (especially, BAC end sequences, abbr. BES) is shared in a row pool and a column pool, it will be assigned to the well at the intersection of the row pool and the column pool. BAC clones will be further mapped to the reference genome according to assigned contigs.

DNA of the secondary pools were extracted and Illumina sequencing libraries with individual index sequences for each secondary pool were prepared. Finally, 96 row pool libraries were mixed in equal amounts as the library X, and 96 column pool libraries were also mixed in equal amounts as the library Y. The average insert sizes of the library X and Y analyzed by the Agilent 2100 Bioanalyzer (Additional file 1: Fig. S1) were 491 bp and 443 bp, respectively. Sequencing was accomplished using the Illumina HiSeq 2000 sequencing platform with paired-end protocol (PE150). The Library X and Y generated 130.82 Gb and 133.50 Gb raw data, respectively. Trimmomatic was employed for trimming adaptors and filtering low-quality or shorter reads. FastQC was employed for evaluating the quality of the preprocessing reads. Then, the valid reads were obtained by filtering the reads traced to *E. coli* DH10B reference genome using Bowtie2. Finally, we obtained valid yields of 60.40 Gb and 102.37 Gb for libraries X and Y, respectively (Table 1). Demultiplexer was employed for demultiplexing and generating each secondary pool reads according to the 7-bp index (Additional file 2). The average valid sequencing depths of row pools and column pools were 45x and 78x, respectively (Additional file 1: Fig. S2).

**Table 1.** The summary of NGS data.

| Library | Total size (Gb) | Size after QC (Gb) | Valid size (Gb) |
| --- | --- | --- | --- |
| X | 130.82 | 103.78 | 60.40 |
| Y | 133.50 | 107.92 | 102.37 |

### 2.3 Parsing BAC end sequences

We focused our interest on the two special contigs in a BAC: the forward and reverse BAC end seqeunces. We designed two pathways to find them. On one hand, the paired-end reads overlapping with the border sequences of the vector harboring the *Hin*dIII restriction enzyme site were extracted from the valid data of each secondary pool, and then assembled using Cap3 based on the overlap-layout-consensus method. The vector sequence parts in the consensus were truncated to obtain short BAC end sequences (abbr. BES) starting with AAGCTT (*Hin*dIII site). These short BAC end sequences were assigned to the corresponding wells according to the clone array pooled shotgun sequencing strategy. Of the 9,216 wells in 384-plates tested in the pipeline, 8,183 (88.79%) wells were assigned one forward BES and 7,897 (85.69 %) wells were assigned one reverse BES (Additional file 3: Table S1). All assigned BAC end sequences have an average length of 358 bp (Additional file 1: Fig. S3), which is consistent with the insert size of the Illumina libraries (Additional file 1: Fig. S1).

On the other hand, the valid NGS data of each secondary pool were firstly assembled using SPAdes, and the contig N50 sizes of all row pools and column pools were counted (Additional file 1: Fig. S4). The average N50 of row pools and column pools were 13.97 kb and 11.45 kb, respectively. Likewise, the vector sequence parts at the ends of the contigs were removed to retain the long BESs starting with AAGCTT. Finally, these long BESs were assigned to the corresponding wells in 384-plates. Of the 9,216 wells, 5,454 (59.18%) wells obtained one forward BES and 5,108 (55.43%) wells obtained one reverse BES (Additional file 3: Table S1). The N50 of the long BESs is 60.87 kb (Additional file 1: Fig. S5).

### 2.4 Determination of BAC locations on the genome

In order to determine the locations of the broomcorn millet BAC clones on the genome, we aligned the short and long BAC end sequences onto the cultivar longmi4 genome with Blastn. Table 2 listed the alignment results. With BAC end sequences with repeats, 14,862 (83.44%) short BAC end sequences including 12,971 single-hit and 1,891 multi-hit sequences were mapped to the genome, and 12,120 (95.28%) long BAC end sequences (all single-hit) were mapped to the genome. With the BAC end sequences without repeats, 7,760 (43.57%) short BAC end sequences including 7,295 single-hit and 465 multi-hit sequences were mapped to the genome, and 10,626 (83.53%) long BAC end sequences (all single-hit) were mapped to the genome. In order to map as many as possible BACs to the genome, the alignment results of the end sequences with repeats were adopted. We wrote a python script to extract the Blast results from short and long BESs. As a result, 5,795 BACs were mapped to the genome using short BAC end sequences, and 6,973 BACs were mapped to the genome using long BAC end sequences. Finally, the Blast results generated by short and long BAC end sequences were integrated, and in total 8,262 BACs (89.65%) were mapped to the genome.

**Table 2.** A summary of the Broomcorn millet BESs and the anchoring results of the Broomcorn millet BAC clones to the longmi 4 genome using the BESs

| Categories | Short BESs | Long BESs | Integrity |
| --- | --- | --- | --- |
| BAC end sequences |  |  |  |
| Clones in superpool | 9216 | 9216 |  |
| Clones with successful BESs | 9126 | 7725 |  |
| with paired BESs | 7790 | 3890 |  |
| with single-end BESs | 1336 | 3835 |  |
| only forward | 921 | 2189 |  |
| only reverse | 415 | 1646 |  |
| Total successful BESs | 17811 (96.63%) | 12721 (66.17%) |  |
| Alignment |  |  |  |
| Aligned BESs with repeats unmasked | 14862 (83.44%) | 12120 (95.28%) |  |
| Single-hit BESs | 12971 | 12120 |  |
| Multi-hit BESs | 1891 | 0 |  |
| Aligned BESs with repeats masked | 7760 (43.57%) | 10626 (83.53%) |  |
| Single-hit BESs | 7295 | 10626 |  |
| Multi-hit BESs | 465 | 0 |  |
| Anchoring to reference sequences |  |  |  |
| Clones anchored to single sites | 5795 | 6973 | 8262 |
| Clones anchored with single BES | 2507 | 3907 | 2871 |
| Clones anchored with paired BESs | 3288 | 3066 | 5391 |

To verify the accuracy of BAC locations on the genome, 55 BAC clones were randomly picked from 384-plates for BAC end sequencing using Sanger method. After quality control, the 55 paired Sanger end sequences were blasted with the above short and long BESs, and the genome. All BAC clones but one (32G16) were consistent. By checking the 32G16 BAC end sequences, we found that this clone (or rather 32G16 well) was assigned two long forward BESs, one short forward BES, and no reverse BES. Only one long forward BES and the short forward BES were perfectly identical to the Sanger end sequence. Another long forward BES confused the mapping. However, it can be solved by weighting. In summary, the accuracy of the BAC mapping approach in this study was extremely high.

The distribution of the 5,391 BACs mapped to the genome by paired BAC end sequences were counted (Table 3). These clones covered a total of 432.47 Mb of chromosomes with a total coverage of 50.97%. Among the 18 chromosome sequences of the cultivar longmi4 genome, there left 829 gaps. Our BACs covered 308 of them. The insert sizes of these BACs presented the Gaussian distribution, with an average insert size of 123.48 kb (Fig. 3), which is lower than that predicted by pulse field electrophoresis. These BACs are valuable resources for further improvement of the genome.

**Fig. 3.**
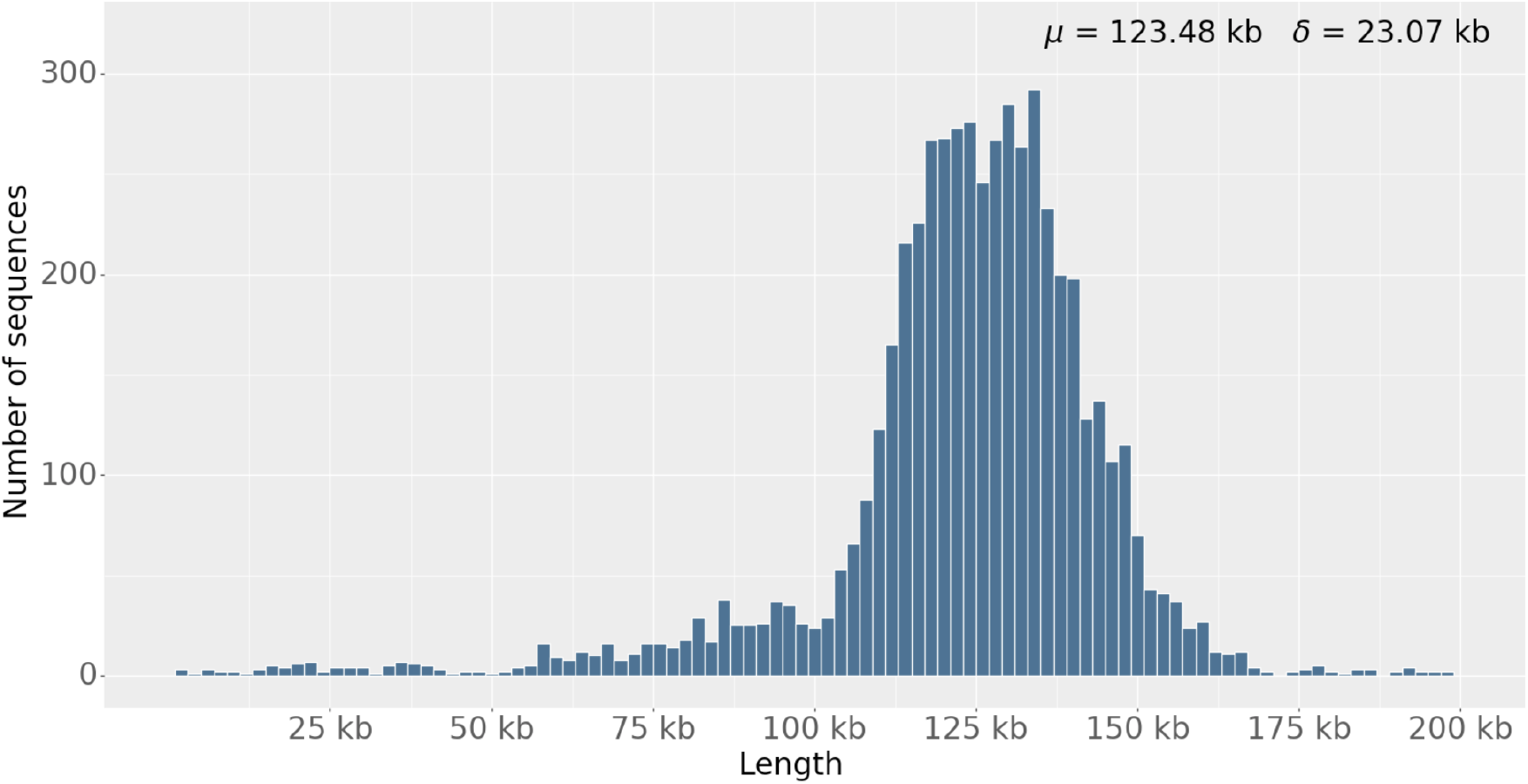
the statistics of the BAC inserts mapped by paired BESs

**Table 3.** The location result of BACs on broomcorn millet chromosomes

| Chr. | Length (Mb) | No. gaps | No. BACs | BACs Covered |  |
| --- | --- | --- | --- | --- | --- |
|  |  |  |  | length (Mb) | No. gaps |
| chr 1 | 69.18 | 85 | 439 | 35.25 | 34 |
| chr 2 | 61.15 | 59 | 379 | 31.04 | 19 |
| chr 3 | 57.97 | 50 | 394 | 31.03 | 20 |
| chr 4 | 56.29 | 34 | 359 | 27.90 | 14 |
| chr 5 | 54.13 | 52 | 361 | 29.86 | 19 |
| chr 6 | 52.84 | 46 | 358 | 28.24 | 16 |
| chr 7 | 51.23 | 67 | 286 | 23.67 | 27 |
| chr 8 | 48.26 | 29 | 339 | 26.49 | 8 |
| chr 9 | 45.11 | 70 | 240 | 20.96 | 21 |
| chr10 | 44.65 | 53 | 308 | 25.44 | 28 |
| chr11 | 43.18 | 30 | 267 | 21.90 | 14 |
| chr12 | 42.47 | 30 | 254 | 20.79 | 11 |
| chr13 | 40.72 | 50 | 261 | 21.11 | 17 |
| chr14 | 38.49 | 32 | 269 | 20.23 | 10 |
| chr15 | 34.36 | 34 | 213 | 18.14 | 8 |
| chr16 | 33.61 | 45 | 212 | 16.72 | 15 |
| chr17 | 32.99 | 25 | 226 | 18.26 | 10 |
| chr18 | 32.24 | 38 | 199 | 15.32 | 17 |
| unplaced | 9.48 | 0 | 27 | - | 0 |
| Total | 848.47 | 829 | 5391 | 432.47 | 308 |

### 2.5 Presentation of BAC locations by JBrowse

In order to effectively use the BAC resource, quickly and easily retrieve BAC clones and view other annotaion information, we established a resource website employing a lightweight JBrowse to display the BAC resource information on the broomcorn millet genome (Fig. 4).

**Fig. 4.**
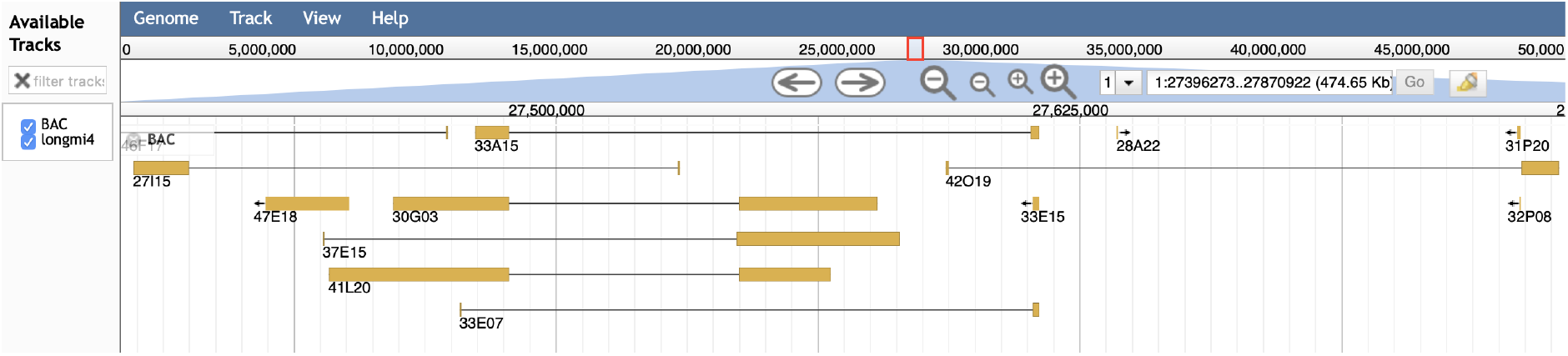
Presentation of broomcorn millet BAC locations by JBrowse. A barbell icon idicates a BAC mapped by forward and reverse BESs (yellow rectangle). An icon with arrow indicates a BAC mapped by only single BES.

### 2.6 Organelle genomes of the broomcorn millet

We aligned all BESs in this study to the chloroplast (CM009689) genome of broomcorn millet by local blast. A total of 65 BAC clones were mapped to the chloroplast with more than 99% of similarity, of which 16 BAC clones were determined by paired BESs (25C07, 25N09, 27F16, 27O19, 29B02, 29K11, 32J09, 32N16, 33D23, 33K24, 36A05, 39C22, 42C10, 46C16, 48B15, 48C03). In the absence of mitochrondrial sequence in broomcorn millet, we aligned BESs to all mitochondrial genomes in *Gramineae*. Of 16 BAC clones with homologous BES sequences, 13 BACs were simultaneously aligned to the chloroplast and 3 BACs were simultaneously aligned to the nuclear genome.

## 3 Discussion

BAC library is still a powerful resource for genome assembly, functional genomics research, and long-term genetic resoure storage for the endangered species. BAC seqeunces, espcially paired BESs, are usally used to detect assembly errors or assist assembly [12]. The most conventional and convenient approach to decode BESs is Sanger sequencing method. However, its one-by-one style is laborious, time-consuming and expensive. Fortunately, a few of efficient approaches had developped, based on next generation sequencing technologies, such as pBACcode and BAC-anchor[13, 14]. pBACode determines paired BESs by a pair of random barcodes flanking the cloning site. BAC-anchor determines general paired BESs by using specific restriction enzyme sites and searching for utlra-long paired-end subreads containing large internal gaps. We previously also developed a high-throughput approach for long accurate BES profiling with PacBio sequencing technology [15]. However, these approaches focused on paired BES profiles, and they are hard to trace BESs of specific clones in 384-plate wells. In this study, we generated BESs and assigned them to physical wells in 384-plates by applying the characteristics of row-cross-column in two-dimension arrays and cost-effective Illumina platform. These BAC end sequences in average are longer and accurater than that generated by Sanger method, for assembling by SPAdes assembler[16]. After alignment, 89.65% of clones were successfully associated with the broomcorn millet longmi 4 genome. These clones covered 308 of the 829 gaps left by the genome and can be uesd to close the genome gaps.

In conventional whole genome sequencing project, sequencing coverage is usallly at least 30x, and not more than 200x. Higher coverage will result in more sequences that are generated by PCR mutations or sequencing errors and lead to contigs with shorter N50. We added index into each secondary pool and mixed 96 secondary pools as a sequencing library to run in a lane of Illumina flow cell for greatly reducing the cost of sequencing. As a result, after removing the contaminated *E. coli* genomic DNA reads the average valid sequencing depths of row pools and column pools were 45x and 78x, respectively.

In the process of BESs extraction, we designed two pathways: short BES pathway and long BES pathway. Short BES pathway searched all reads overhanging with verctor end sequences before assembling by Cap3; consequently, it generated all potenital BESs. Long BES pathway identified all contigs overhanging with vector end seqeunces after assembling by SPAdes; consequently, it generated longer but less BESs than the first pathway. The assignment of BAC end sequences at the intersection sites is affected by many factors, such as the overlapping rate of the BAC clones and the correct rate of the sequences. The overlapping rate of BAC clones in the superpool is the most important factor. If overlapping BAC clones appear in the same row or column pool, we can assign them to wells easily and correctly. Also If two overlapping BAC clones appear in a different row or column pool, we can rectify them by our previous method [17]. However, if more than two overlapping BAC clones appear in a different row or column pool, our method will filter out potential BESs, so that the BESs of the intersection well will be absent. In the process of BES assignment, the flow of the forward and reverse BESs are completely independent. When more than one forward and/or reverse BESs were assigned to a well, we cannot determine which BESs are a pair of BESs. If a high-qualty genome is available, it is easy to assess which pair of BESs are derived from the same BAC by mapping. However, if the variety used for BAC library construction is not the same as that the reference genome stands for, the alignment results that do not satisfy the location requirement of BES pairs will be discarded. The clones that may contain a large structural variation can be picked out for further analysis from the 384-plate wells.

Plant cells contain an abundance of chloroplasts and mitochondria. The chloroplast genome is generally around 150 kb, while the mitochondrial genome size varies widely, typically between 200 kb and 750 kb [18]. Although nuclei are extracted for BAC library construction, a trace of organelle DNA cotamination is inevitable. In this study, a small number of clones of chloroplast genome was found by BESs, while clones of mitochondrial genome were almost absent.

By high-throughput sequencing and mapping clones in the secondary pool of BAC libraries, we can find the coordinates of genome sequences in the broomcorn millet BAC libraries. Therefore, if we find genes that play an important role in biology, we can quickly locate the BAC clones containing this gene, and obtain the experimental materials for further research and analysis. At the same time, because BAC contains a long DNA fragment (about 120 kb), it is also convenient for us to quickly analyze the upstream and/or downstream DNA elements of interested genes or adjacent genes.

## 4 Conclusions

We constructed a high-quality BAC library for broomcorn millet, developed a high-efficient and low-cost pipeline which can generate and parse ten thousand of paired BAC end sequences at each time, and mapped a total of 8,262 broomcorn millet BACs to the chromosomes. These clones covered 308 of the 829 gaps left by the genome. The high-quality BAC clones mapped on genome in this study will provide a powerful genomic resource for gap filling, complex segment sequencing, FISH, functional research, and genetic engineering of broomcorn millet.

## 5 Methods

### 5.1 Plant materials, growth conditions and BAC library construction

The seeds of broomcorn millet were provided by professor Mingsheng Chen of the Institute of Genetics and Developmental Biology, Chinese Academy of Sciences and grown at dark conditions under 25°C. The young leaves of seedlings were harvested and mixed, frozen immediately in liquid nitrogen for the extraction of nuclear DNA. High molecular weight nuclear DNA was extracted and BAC library was constructed following our previous protocol [6]. Partial digestions of DNA plugs with dilution *Hin*dIII were performed. DNA fragments ranging from 100 kb to 200 kb were recovered from pulse field gel and ligated with pIndigoBAC536-S vector [7]. The ligation product was used to transform DH10B cells by electroporation. White colonies were picked up and stored in 384-plates at −80 °C.

### 5.2 Pool construction

Twenty-four 384-plates were chosen form the BAC library of broomcorn millet. These plates were arranged in a 2-dimension superpool. The superpool contains 96 row pools and 96 column pools. A total of 192 pools were processed independently for high quality DNA extraction with the AxyGen AxyPrep Easy-96 plasmid kit. Then, these BAC DNAs were completely digested with ATP-Dependent DNase (Epicentre) to remove the host *E. coli* DNA. After digestion, these BAC DNAs were sheared in the Bioruptor to an average of 500 bp. During blunt-end repair, overhanging 5’ and 3’ ends were filled in or removed by T4 DNA polymerase. 5’-phosphates were attached using T4 polynucleotide kinase. Tail-A were added using Taq DNA polymerase. Then, adapters were ligated to both ends of the molecules using T4 DNA ligase. The ligation products were cleaned using MagBead DNA Purification Kit (Sangon Biotech, shanghai, CN). Sequencing adaptors were added using PCR. Finally, the products of KOD PCR were cleaned again.

### 5.3 Illumina sequencing and BAC end sequence analysis

NGS sequencing of 2 mixed DNA libraries was performed via the Illumina HiSeq 2000 with 150-bp paired-end protocol (Genewiz, suzhou, CN). Trimmotatic was employed to filter and trim raw reads. FastQC was employed to assess data quality. After adapter filtering and quality assessment, BBMAP/demuxbyname script was employed for deconvolution depending on unique index sequence in each pool. A python script was employed to extract target paired-end reads that cover BES-VES site. Cap3 was employed to assemble those reads to consensuses, and consensuses were then trimmed to remove the part of vector to generate BESs called “short BES”. SPAdes was employed to directly assemble pool reads to contigs. Then, the contigs containing BAC vector sequences were extracted and trimmed using python script to generate the trimmed contigs called “long BES”. Blastn was used to align BESs from raw and column pools, and then the shared BESs were assigned to the wells at the intersection.

### 5.4 Validation of BAC end sequences

The analysis results of the BAC end sequences were validated by Sanger sequencing. Fifty-five BAC clones were randomly selected and their DNAs were extracted using an improved alkaline lysis protocol. Sanger sequencing was accomplished using BAC-F (5’-AACGACGGCCAGTGAATTG-3’) and BAC-R (5’-GATAACAATTTCACACAGG-3’) primers from pIndigoBAC536-S vector backbone. The BAC end sequences from Sanger sequencing were aligned to BES from the analysis results of illumina data using Blastn.

### 5.5 BAC Mapping on broomcorn millet longmi4 genome

The genome sequence of broomcorn millet longmi4 was downloaded from NCBI genome database under GCA_002895445 accession, which was submitted by researchers from China Agricultural University. The local blastn was employed to map all BESs to the genome with the following options: qcov_hsp_perc=99, perc_identity=99, outfmt=6, culling_limit=1. The results were further converted to a GFF3 format file using a Python script. In this script, following conditions were set: if both forward and reverse BESs in each clone were mapped to the same chromosome, and their orientations were opposite and their interval lengths were less than 250 kb, such clones were recorded in GFF3 format file with three lines; if either forward or reverse BES was uniquely mapped to chromosome, such clones also were recorded in GFF3 format file with two lines; other conditions would be discarded. The GFF3 file was sorted using GFF3sort and presented with JBrowse in our website (http://eightstarsbio.com/gresource/JBrowse-1.16.5/index.html).

### 5.6 Organelle genome analysis

The chloroplast genome (CM009689) of brromcorn millet and all mitochondrial genomes (NC_008331, NC_007982, NC_036024, NC_031164, NC_029816, NC_022714, NC_022666, NC_013816, NC_007886, NC_011033, NC_008362, NC_008360, NC_008332, NC_008333) in *Gramineae* were downloaded form NCBI nucleotide database. All potential BACs of organelle genome were identified by the local blastn with following options: qcov_hsp_perc=99, perc_identity=99, outfmt=6.

## Abbreviations

BAC: bacterial artificial chromosome;
BES: BAC end sequence;
CAPSS: clone array pooled shotgun sequencing;
CM: chloramphenicol;
NGS: next generation sequencing;
PCR: polymerase chain reaction;
VES: vector end sequence.

## 7 Declarations

### 7.1 Ethics approval and consent to participate

Not applicable

### 7.2 Consent for publication

Not applicable

### 7.3 Availability of data and materials

The source codes are openly available in a GitHub repository (https://github.com/xuweixw/broomcorn-millet-BAC-library). Illumina sequencing data are available at Sequence Read Archive (SRA) under the accession PRJNA576359. BAC clones can be browsed and obtained through our website (http://eightstarsbio.com/gresource/JBrowse-1.16.5/index.html).

## 7.6 Authors’ contributions

WX and MLuo conceived and designed the research framework; WX, MLiang, XY and HW performed the experiments; WX analyzed the data and wrote the manuscript; MLuo supervised the work and finalized this manuscript. All authors read and approved the manuscript.

## 7.7 Acknowledgements

We are grateful to Dr. Mingsheng Chen (the Institute of Genetics and Developmental Biology, Chinese Academy of Sciences) for providing the seeds of broomcorn millet.

## 7.8 Authors’ information (optional)

^1^College of Life Science and Technology, Huazhong Agricultural University, Wuhan 430070, China

