## Additional file 1 for "Genomic Resources of Broomcorn Millet: Demonstration and Application of a High-throughput BAC Mapping Pipeline"

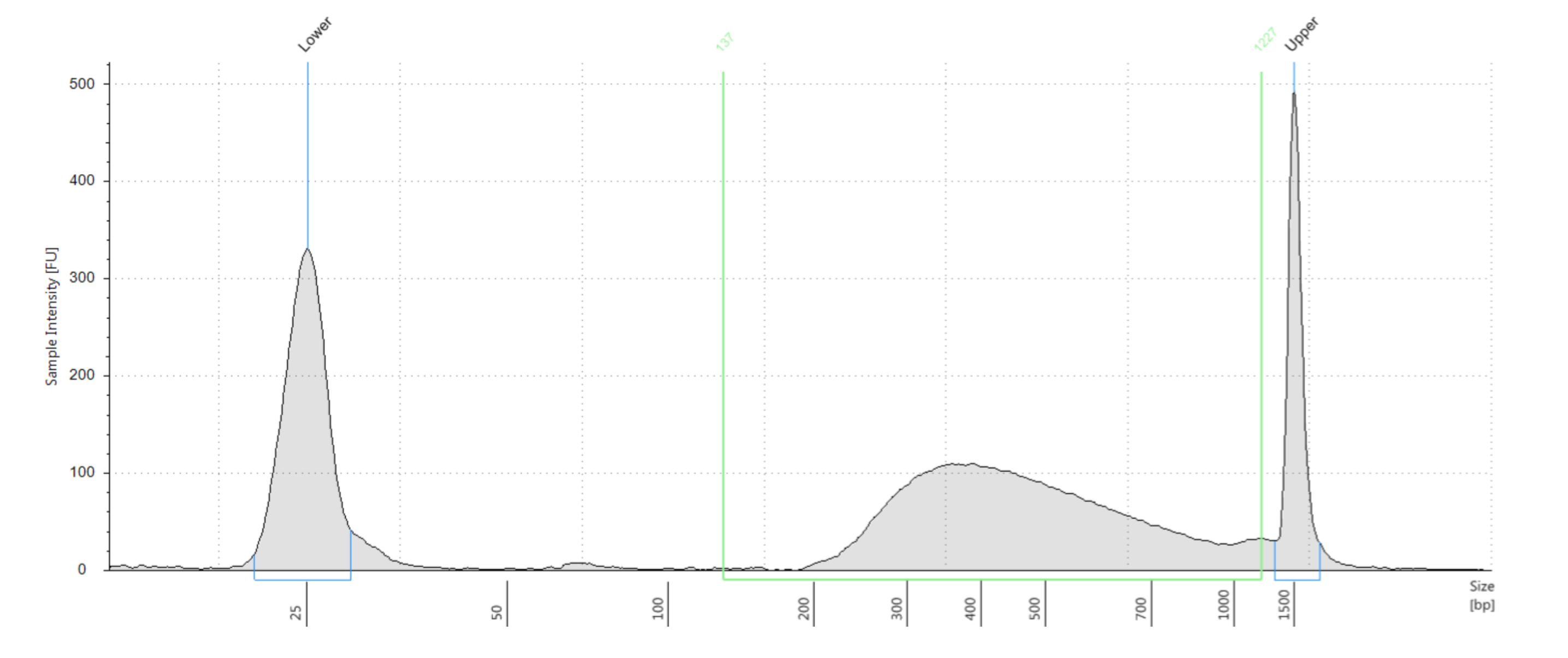

(A)

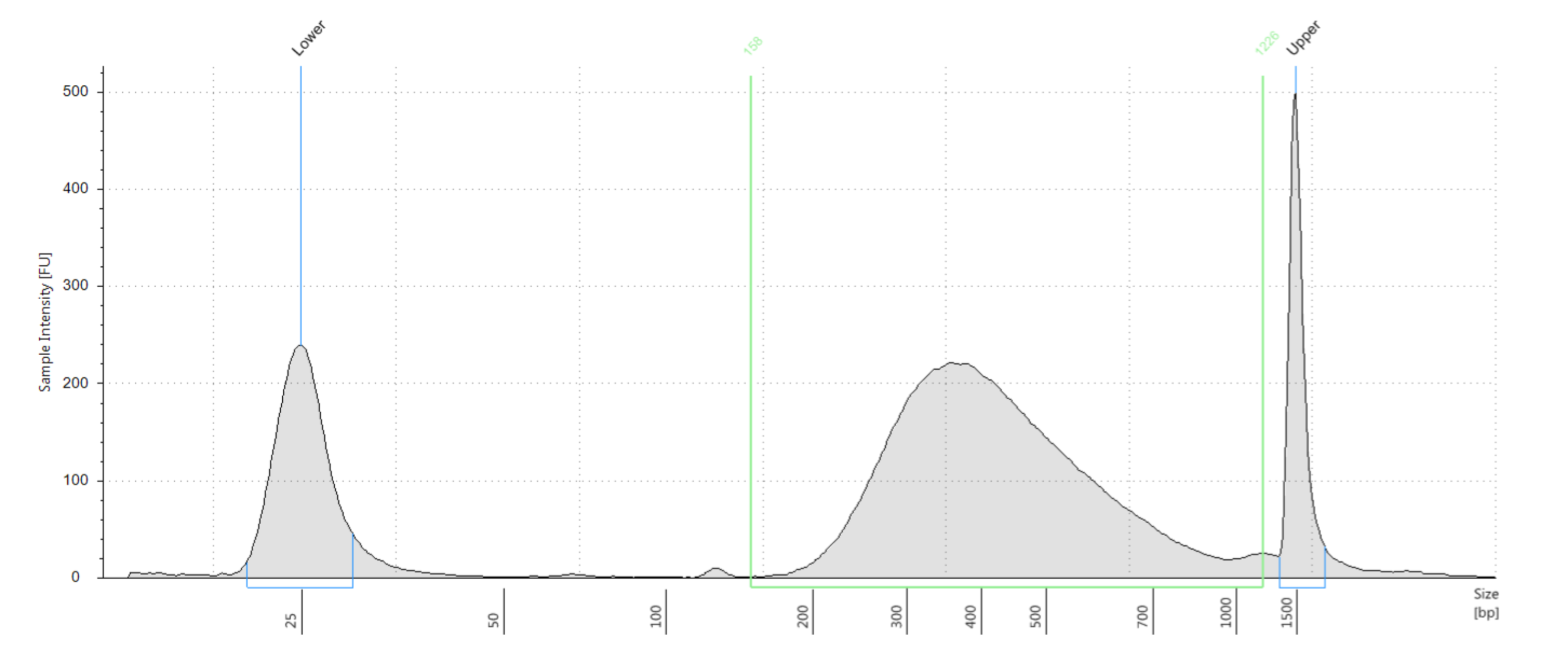

(B)

Fig. S1. Distribution of DNA fragment lengths in library X and Y analyzed by the Agilent 2100 Bioanalyzer. The peak in the middle of each panel is the length range of DNA library. The peaks on the left and right of each panel are 25 bp and 1500 bp DNA standards, respectively. (A) Sequencing library X, (B) Sequencing library Y.

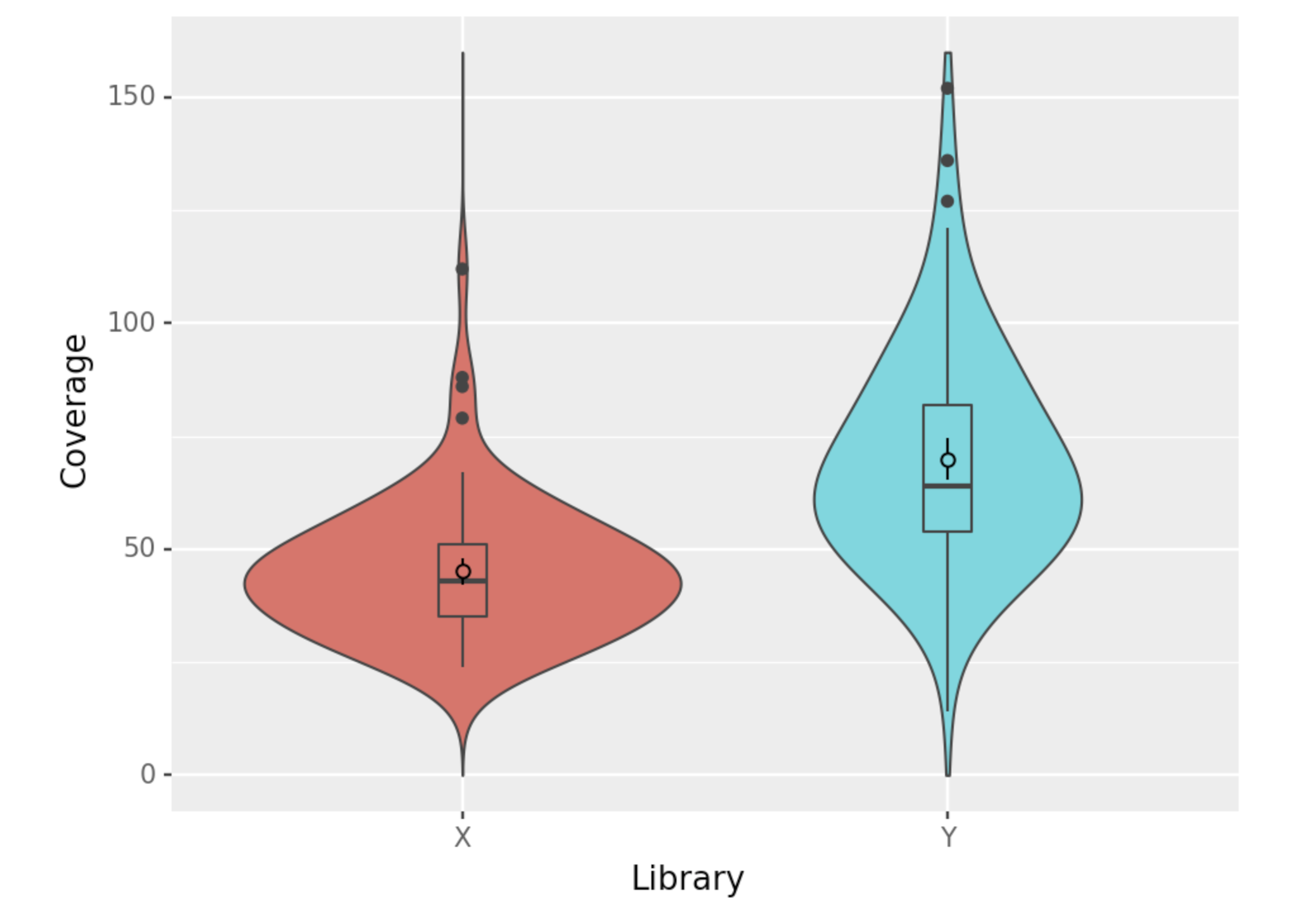

45

78

Fig. S**2** Distribution of the coverage numbers of valid reads in pools of sequencing library X and Y. The average BAC insert size estimated by PFGE is 132 kb and the length of pIndigoBAC536-S vector is 7 kb, so the coverage number of valid reads in each pool is calculated with an average BAC size of 139 kb.

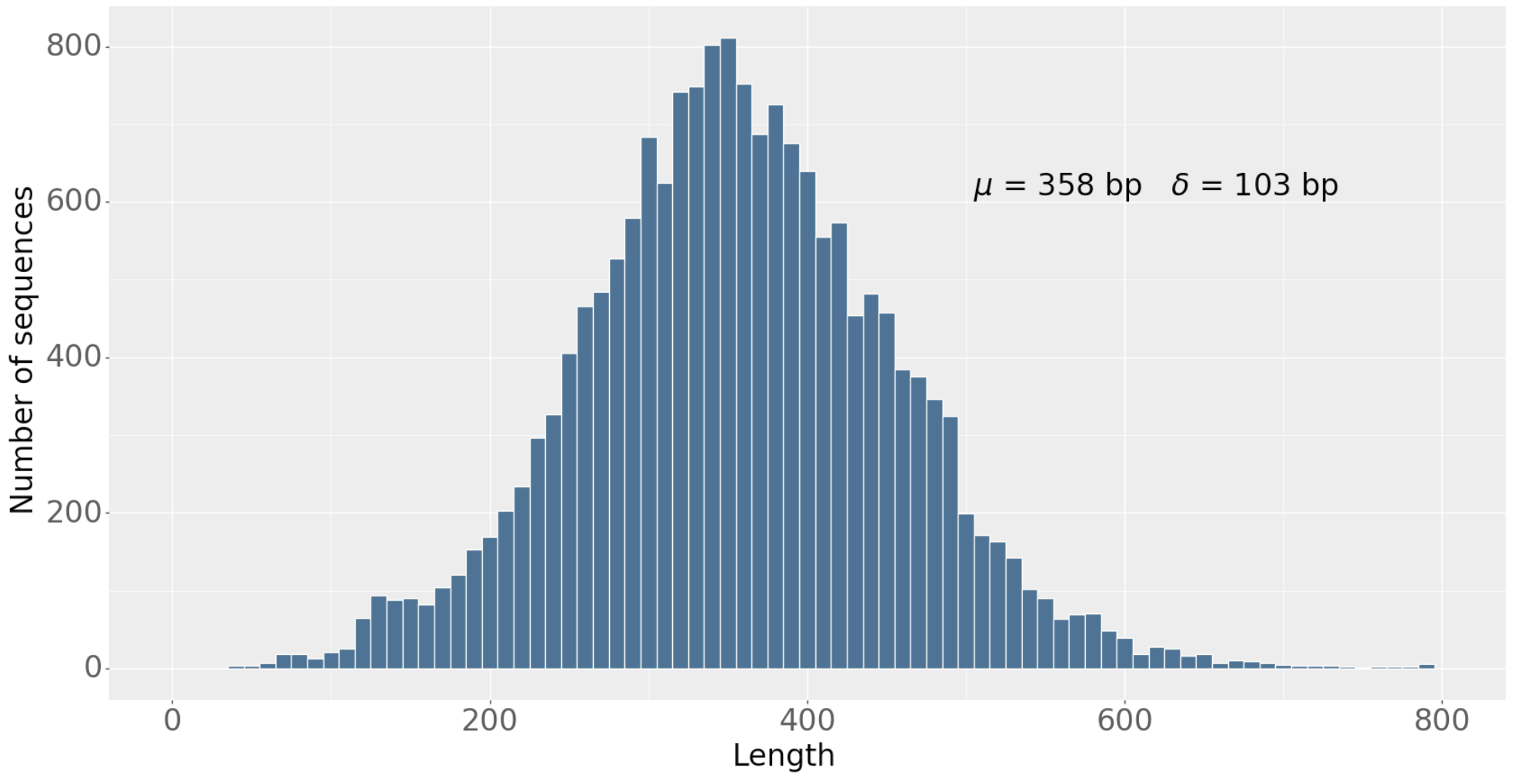

Fig. S3. Distribution of short BAC end sequence lengths

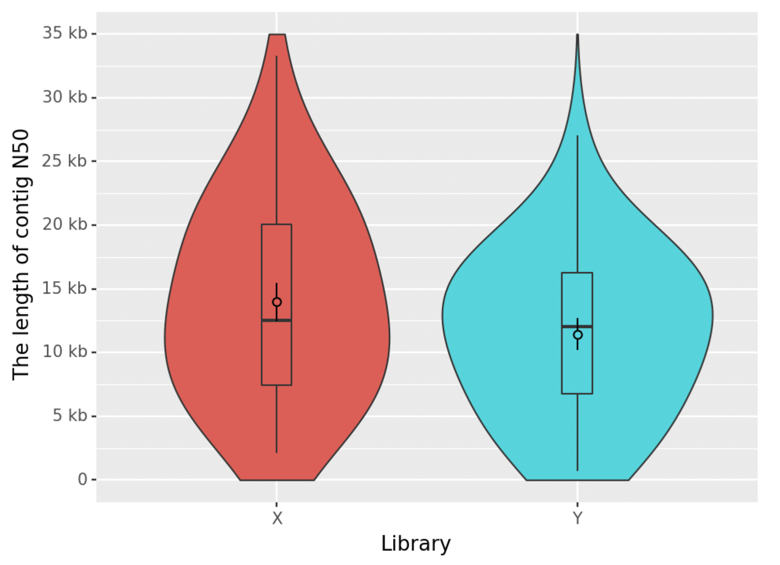

Fig. S4 Distribution of the lengths of contig N50 assembled by SPAdes in pools of sequencing library X and Y.

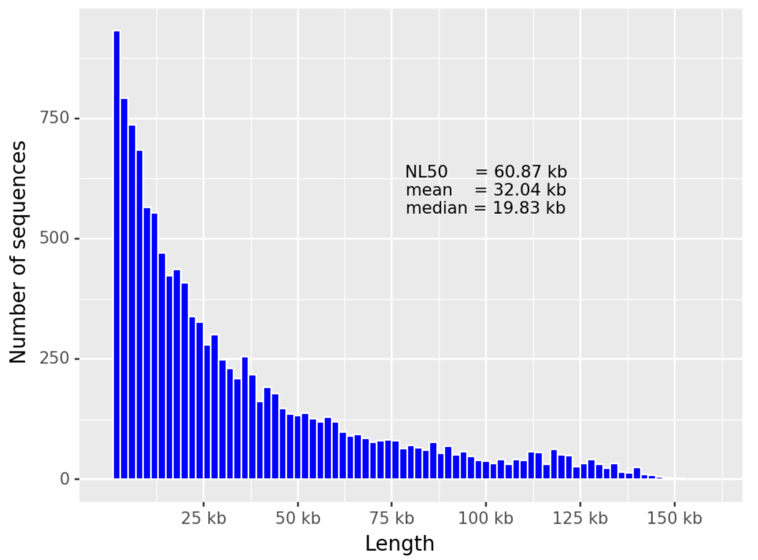

Fig. S**5**. Distribution of long BAC end sequence lengths
