## Additional file 3 for "Genomic Resources of Broomcorn Millet: Demonstration and Application of a High-throughput BAC Mapping Pipeline"

Table S1 the number of wells assigned with BESs

| The Number of BESs | In short BES pathway | | In long BESs pathway | |
| --- | --- | --- | --- | --- |
|  | Forward | Reverse | Forward | Reverse |
| 0 | 505 | 1011 | 3138 | 3680 |
| 1 | 8183 | 7897 | 5454 | 5108 |
| 2 | 487 | 302 | 591 | 410 |
| 3 | 39 | 6 | 31 | 17 |
| 4 | 2 | - | 2 | 1 |
